# Comparative phenotyping of mice reveals canonical and noncanonical physiological functions of TRα and TRβ

**DOI:** 10.1101/2023.11.26.568063

**Authors:** GS Hönes, D Geist, C Wenzek, PT Pfluger, TD Müller, A Aguilar-Pimentel, OV Amarie, L Becker, N Dragano, L Garrett, SM Hölter, B Rathkolb, J Rozman, N Spielmann, I Treise, E Wolf, W Wurst, H Fuchs, V Gailus-Durner, MH de Angelis, D Führer, LC Moeller

## Abstract

Thyroid hormone (TH) effects are mediated through TH receptors (TRs) TRα1, TRβ1, and TRβ2. The TRs bind to thyroid hormone responsive elements on the DNA and regulate expression of TH target genes as ligand dependent transcription factors (canonical signaling). In addition, the TRs α and β mediate activation of signaling pathways, e.g. the PI3K/AKT and MAPK/ERK pathways (noncanonical signaling). Whether such DNA-binding independent TR action contributes to the spectrum of physiological TH effects is largely unknown. The aim of this study was to attribute physiological effects to the two TR isoforms α and β and their canonical and noncanonical signaling. We conducted multi-parameter phenotyping in male and female TR knockout mice (TRα^KO^, TRβ^KO^), mice with disrupted canonical signaling due to a mutation in the TR DNA-binding domain (TRα^GS^, TRβ^GS^) and their respective wild-type littermates. Perturbations in senses, especially hearing (mainly TRβ with a lesser impact of TRα), visual acuity and retinal thickness (TRα and TRβ), in muscle metabolism (TRα) and in heart rate (TRα) highlighted the role of canonical TR action. Strikingly, selective abrogation of canonical TR action often had little to no phenotypic consequence, suggesting that noncanonical TR action sufficed to maintain the wild-type phenotype for specific effects. For instance, macrocytic anemia, reduced retinal vascularization or increased anxiety related behavior were only observed in TRα^KO^, but not TRα^GS^ mice. Noncanonical TRα action increased the efficiency of energy utilization and prevented hyperphagia observed in TRα^KO^ mice. In summary, by examining the phenotypes of TRα and TRβ knockout models alongside their DNA-binding-deficient GS mutants and wildtype counterparts, we could establish that the independent noncanonical actions of TRα and TRβ play a crucial role in modulating sensory, behavioral, and metabolic functions. This comparison underscores the significance of the TRs in orchestrating a spectrum of physiological processes beyond their traditional genomic pathways.

## Introduction

Thyroid hormone (TH; T4, 3,5,3’,5’-tetraiodothyroxine, thyroxine and its active form T3, 3,5,3’-triiodothyronine, thyronine) is essential for development and physiology ^1^. TH effects are mediated through TH receptors (TRs) TRα1, TRβ1, and TRβ2. The TRs bind to thyroid hormone responsive elements (TREs) on the DNA and regulate expression of TH target genes as ligand dependent transcription factors. This TR action by direct binding of TRs to DNA is referred to as canonical or type 1 signaling TH/TR action ^2,3^. In addition, the TRs α and β mediate activation of pathways including phosphatidylinositol-3-Kinase (PI3K)/AKT and MAPK/ERK ^4–8^, which is known as noncanonical or type 3 TH/TR signaling.

To determine TR isoform functions, knockout (KO) mice for TRs α and β have been generated ^9^. Studies of TRα and TRβ KO mice allowed the characterization of TR isoform functions. Predominantly TRα-regulated are bone development and linear growth, maturation of the small intestine and heart rate, whereas TSH suppression and hepatic gene expression are predominantly regulated by TRβ ^10^.

An important question is whether DNA-binding independent noncanonical TR actions contribute to the overall physiological effect of TH. While studies of TR KO mice revealed the contribution of the TR isoforms, they could not determine the mode of TR action underlying the physiological effects, whether canonical or noncanonical, because both modes of action are present in WT mice but absent in TR KO mice. To determine whether DNA binding of TRs is required for physiological effects, we generated mouse models with a mutation in the P box of the DNA-binding domain (DBD), leading to substitution of the first two amino acid (<u>EG</u>G to <u>GS</u>G) of TRα and TRβ were generated (referred to as TRα^GS^ and TRβ^GS^ mice, respectively). *In vitro* EMSA, luciferase reporter assays, ChIP, ChIP-Seq and microarray studies from mouse liver samples confirmed the loss of DNA binding, histone acetylation and transcriptional activity of TRβ due to the GS mutation ^11–13^. This mutation specifically abrogates DNA-binding to TREs while preserving noncanonical TR action. This approach allows us to discern the necessity of DNA binding in eliciting physiological effects, providing valuable insights into the intricate mechanisms of TR-mediated actions. Comparing WT, TR KO, and TR DNA-binding-deficient GS mice clarified each TR isoform’s role in thyroid hormone action and the requirement of DNA binding for their physiological effects: Arterial vasodilation was identified as a rapid noncanonical TRα effect, whereas longitudinal growth and bone development were regulated by canonical TRα action ^13,14^. Negative feedback in the hypothalamus-pituitary-thyroid axis depended on DNA binding of TRβ, whereas liver triglyceride content and rapid effects on blood glucose appeared to be regulated by noncanonical TRβ action ^13^. This prompted us to undertake a much broader approach and assess the contribution of canonical and noncanonical TR signaling to multiple physiological parameters. Specifically, we compared male and female TRα^KO^, TRα^GS^, TRβ^KO^, and TRβ^GS^ mice with their respective WT littermates in a comprehensive multi-parameter phenotyping screen ranging from clinical chemistry and energy metabolism to eye and ear function and behavioral tests, with the aim to attribute physiological effects to the two TR isoforms α and β and their canonical and noncanonical signaling.

## Material and Methods

### Animal breeding and housing

All animal experiments at the University Hospital Essen were approved by the Landesamt für Natur, Umwelt und Verbraucherschutz Nordrhein-Westfalen (LANUV-NRW) and all tests performed at the GMC were approved by the responsible authority (Regierung von Oberbayern). Studies were performed in accordance with the German regulations for Laboratory Animal Science (GVSOLAS) and the European Health Law of the Federation of Laboratory Animal Science Associations (FELASA) (84-02.04.2017.A157 and 84-02.04.2014.A092). TRα KO (TRα^0^) and TRβ KO (TRβ^-^) mice ^9,15^ mice, here referred to as TRα^KO^ and TRβ^KO^, respectively, were acquired from the European Mouse Mutant Archive (https://www.infrafrontier.eu). TRα^GS^ and TRβ^GS^ mutant mice were created with a zinc finger nuclease approach as described previously ^13^. Mice were bred and housed in the central animal facility at the University Hospital Essen. All mice were housed in individually ventilated cages (IVC) at 21±1 °C or 22±1 °C (GMC) in an alternating 12:12 hour light-dark cycle with standard chow and tap water provided according to University Hospital Essen and GMC standard housing conditions (https://www.mouseclinic.de/about-gmc/mouse-husbandry/index.html), respectively. Mice were bred at the same time and subsequently marked with ear punches. They were genotyped at the age of 3-4 weeks and sent to the GMC at the age of 6 weeks *via* overnight shipping. Upon arrival they were housed for 2 weeks before they entered the phenotyping pipeline. Male and female mice were compared which allowed to detect not only canonical and noncanonical TR effects, but also sex-related differences as well.

### Mouse phenotyping pipeline

At the German Mouse Clinic female and male TRα^KO^, TRα^GS^, TRβ^KO^ and TRβ^GS^ mouse cohorts and corresponding wild type controls were subjected to an extensive phenotypic screening in the areas of behavior, neurology, energy metabolism, cardiology, eye, clinical chemistry, immunology and pathology ^16,17^ (www.mouseclinic.de).

A cohort of 10 male and 10 female TRβ^KO^ was compared with an equivalent number of control males and females. Similarly, 10 male and 10 female TRβ^GS^ were compared to the same number of control males and females. Additionally, 12 male and 12 female TRα^KO^ were compared to 15 male and female controls, and 15 male and 15 female TRα^GS^ compared to 14 males and 15 female controls. The phenotyping measurements were taken from weeks 8 to 16 for TRβ^KO^ and TRβ^GS^, and weeks 9 to 19 for TRα^KO^ and TRα^GS^. Assignment of experimental groups was based on the genotype of the animals, and metadata for each data point were recorded throughout the measurements.

The phenotypic tests were part of the GMC screening pipeline and performed according to standardized protocols as described before ^18–23^. Animal numbers may vary depending on the test performed, as indicated in the respective figure or table.

### Statistical analysis

Data were analyzed with GraphPad Prism 8 (GraphPad, San Diego, USA). Detailed information on the statistical analysis can be found in the respective figure legend. Differences were considered significant when P<0.05.

### Data availability

Relevant information about phenotyping data can be requested directly from the corresponding authors.

## Results and Discussion

### Senses

#### Hearing

Compared to their WT littermates, auditory brainstem responses (ABR) of TRα^GS^ and TRα^KO^ mice showed increased sound pressure thresholds at higher frequencies, indicating reduced hearing sensitivity due to the loss of canonical TRα action (Fig. 1A). Compared to TRα mutant mice, both TRβ^KO^ and TRβ^GS^ mice showed a much more severe hearing loss across the entire frequency spectrum with no difference between the two mutant strains, suggesting that normal hearing requires canonical TRβ action (Fig. 1B).

**Figure 1:**
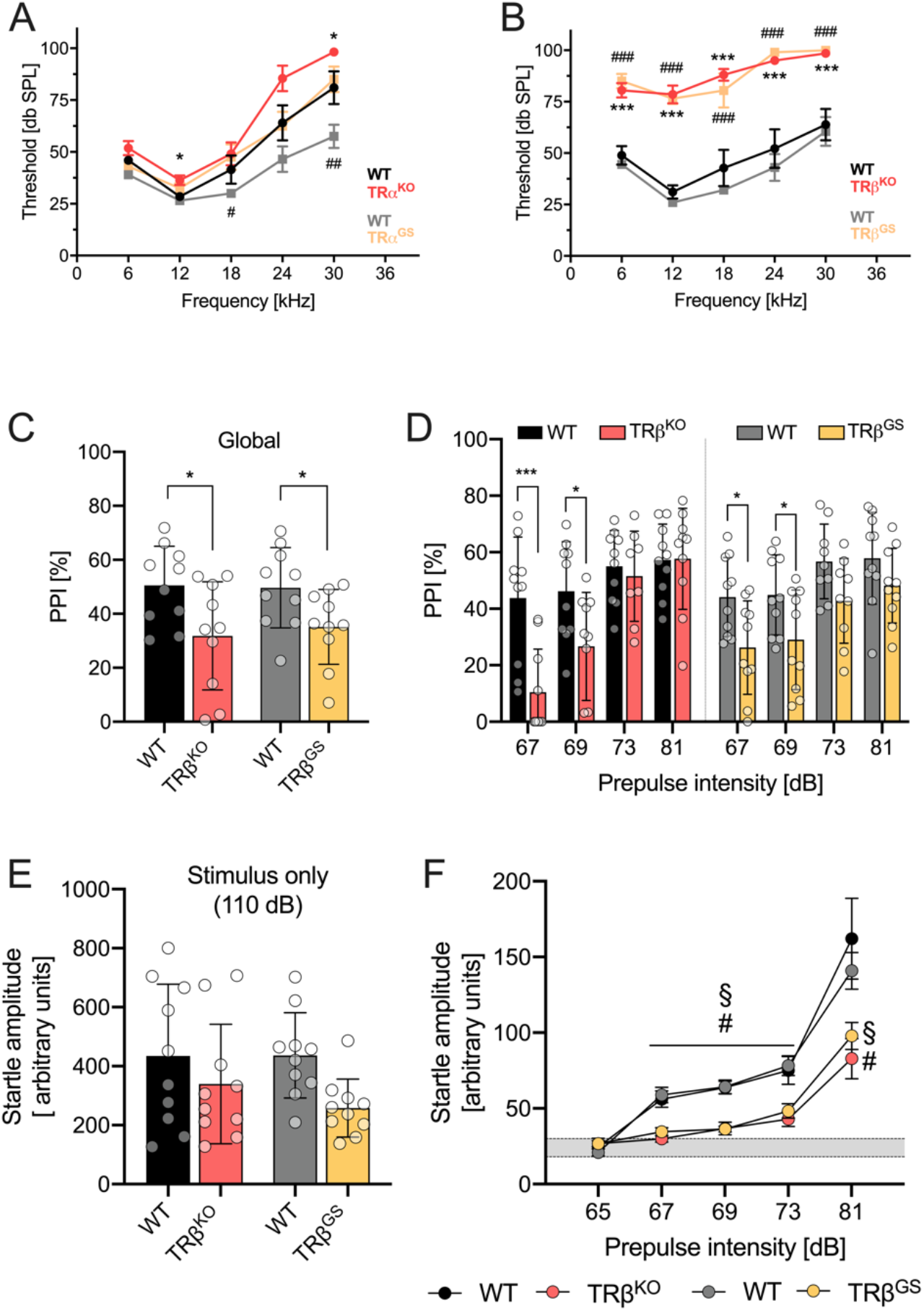
Auditory brainstem responses (ABR) and prepulse inhibition (PPI). ABR of female **A** TRα mutant and **B** TRβ mutant mice. Multiple t-tests; significance: WT vs. TRα^KO^ and WT vs. TRβ^KO^ = \**P<0.05*, \*\*\**P<0.001*; WT vs. TRα^GS^ and WT vs. TRβ^GS^ = ^#^*P<0.05*, ^##^*P<0.01*, ^###^*P<0.001*. **C** global PPI, calculated as the mean PPI [%] for the different prepulse responses. **D** PPI separated by prepulse intensities ranging from 67 to 81 dB. **E** Startle amplitude with a stimulus of 110 dB. **F** Prepulse-alone trials to assess the acoustic startle amplitude on the prepulse *per se*. Multiple t-tests; \**P<0.05*, \*\**P<0.01*, \*\*\**P<0.001;* WT vs. TRβ^KO^ = ^#^*P<0.01*; WT vs. TRβ^GS^ = ^§^*P<0.01.* (TRβ, n= 10) (TRα, n=12-15).

The acoustic startle response and prepulse inhibition (PPI) of both TRα mutant genotypes did not differ from their WT littermates (Fig. S1). Global PPI, calculated as the mean PPI [%] for the different prepulse responses, was decreased in TRβ^KO^ and TRβ^GS^ mice (Fig. 1C). Both TRβ mutant lines showed significantly reduced PPI at intensities of 67 dB and 69 dB but reached levels similar to their WT controls at 73 dB and 81 dB (Fig. 1D). The startle amplitude with a stimulus of 110 dB was slightly but not significantly reduced in TRβ^KO^ and TRβ^GS^ mice (Fig. 1E), whereas prepulse-alone trials confirmed a generally lower startle amplitude in TRβ^KO^ and TRβ^GS^ mice (Fig. 1F). The reduced responses to prepulses up to 69 dB and the reduced startle amplitude up to 81 dB in TRβ mutant mice, along with the absence of differences between TRα mutant mice and WT mice, is consistent with the possibility that the apparent impaired sensorimotor gating in this case reflects the hearing impairment in the mutant mice compared to TRα mutant mice.

Similar results had been previously found in TRα and TRβ KO mice, supporting the conclusion that TRβ1 serves as the major mediator of TH action in the ear, while TRα action is also necessary for complete auditory function ^24–28^. It was suggested that patients with RTHα (resistance to thyroid hormone α) may also suffer from mild hearing loss and the present data support this hypothesis. We did not detect differences between TRβ^KO^ and TRβ^GS^ mice at 15 weeks of age although less pronounced hearing loss had been reported in 8-week-old TRβ^GS^ mice ^12^ and cannot exclude the possibility of progressive hearing loss between weeks 8 and 15.

#### Vision

Posterior eye morphology and fundus development revealed no genotype differences. All retinal layers were present, and lenses and corneas were clear in all genotypes. But retinal thickness was decreased in female TRα (TRα^KO^ and TRα^GS^) and TRβ mutants (TRβ^KO^ and TRβ^GS^) (Fig. 2A). Compared to WT mice, the number of main fundus vessels was decreased in male and female TRα^KO^ but not in TRα^GS^ mice (Fig. 2B). This similarity between TRα^GS^ and WT mice suggests that noncanonical TRα action contributes to retinal vascularization. The spatial frequency threshold, determined with a virtual optokinetic drum as a measure of visual acuity, was decreased in TRα^KO^ mice whereas TRα^GS^ mice were less affected (Fig 2C, left panel). Similarly, this phenotype was observed in TRβ^KO^ mice but not in TRβ^GS^ mice (Fig. 2C, right panel).

**Figure 2:**
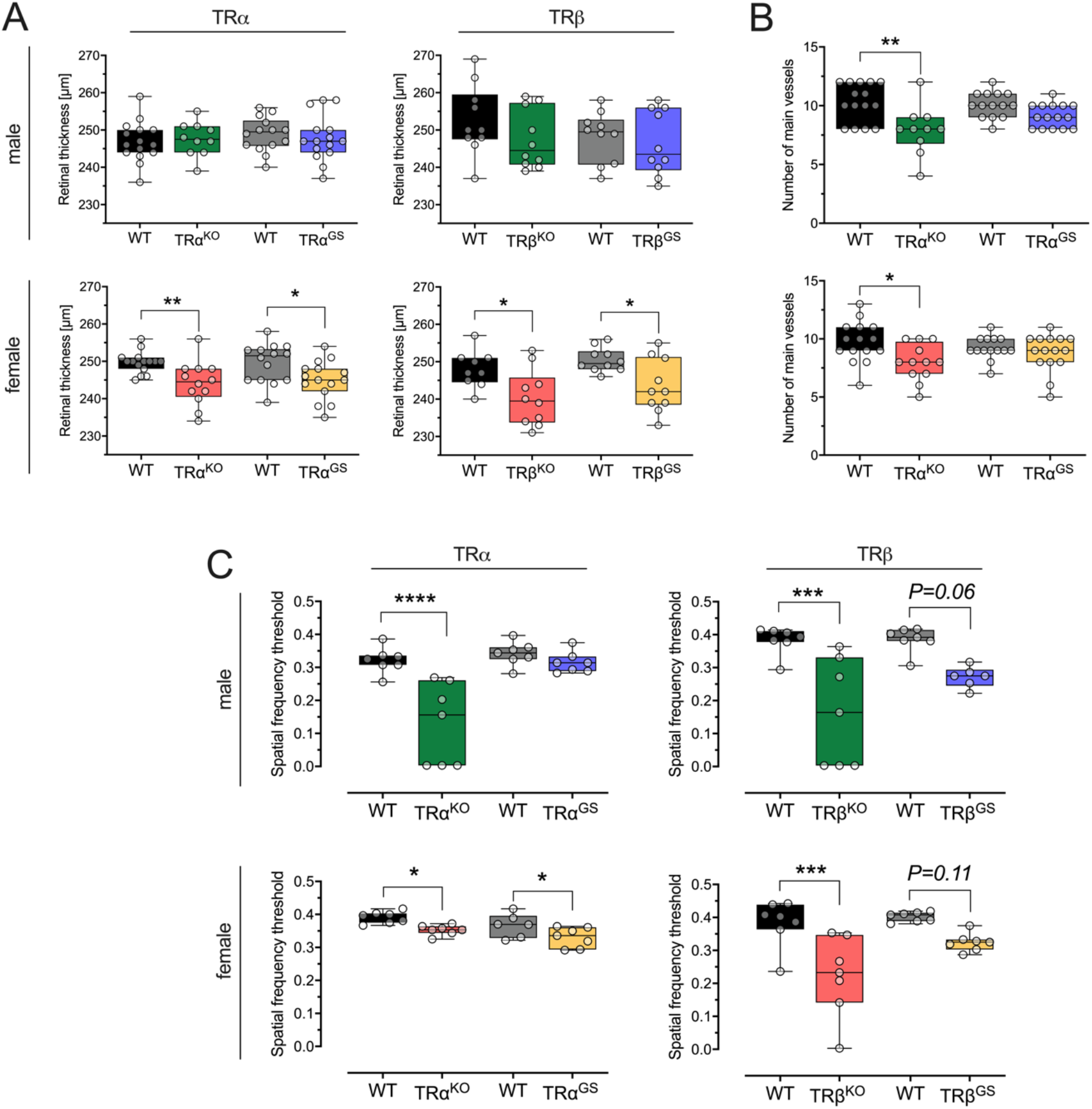
Retinal morphology assessed by optical coherence tomography (OCT) and virtual vision test. **A** Retinal thickness measured by OCT of male and female TRα and TRβ WT and mutant mice. **B** Number of main blood vessels in male TRα mutant mice and WT controls. **C** Visual acuity measured by virtual optokinetic drum (n=7/group/sex). (One-way ANOVA with Sidak’s multiple comparison test; \**P<0.05*, \*\**P<0.01*, \*\*\**P<0.001*, \*\*\*\**P<0.000*1; n=11-15 TRα strains; n=10 TRβ strains).

The impairment of visual acuity and retinal development in TRβ^KO^ mice is expected. TRβ2 regulates cone maturation and the expression of the light-sensing pigment protein cone opsin ^29,30^. In a previous study of TRβ^GS^ and TRβ^KO^ mice, both showed S opsin expression in significantly more dorsal regions compared to WT mice, along with an absence of M opsin expression throughout the retina, which demonstrated that the DNA-binding function of TRβ is essential for cone opsin gene expression in the retina ^12^. The absence of canonical TRβ action appears to impair the development and maturation of the retina, resulting in reduced thickness, which together may contribute to the impaired visual function. Furthermore, female hypothyroid rats studied between embryonal day E19 and postnatal day P5 demonstrated reduced global and layer-specific retinal thickness, neuroblast size and density, ganglion cell density, and mitotic index ^31^, which appears to correspond to the reduced thickness observed here, especially in female TRα and TRβ KO and GS mice.

The *Thra* gene is also expressed in the mouse retina ^32^, but its function remains unknown. The reduced retinal thickness and visual acuity observed in TRα^KO^ mice, which closely resemble the phenotype of TRβ^KO^ mice, indicate that TRα is necessary for normal retinal development and that its absence cannot be compensated by TRβ. The coordinated expression and action of both TRα and TRβ during development could be required for the full maturation of the retina. Additionally, noncanonical TRα action appears to contribute to retinal vascularization. These results suggest that both TRs are necessary for normal retinal development and visual acuity. Further investigation is needed to determine the timing and cell types of the respective TR expressions.

### Behavior

Locomotive behavior, the willingness to explore and anxiety were assessed with the open field test. There were no significant differences in distance travelled for both male TRα mutant mice compared to WT littermates. But rearing activity was markedly reduced in TRα^KO^ mice while TRα^GS^ mice did not differ from WT controls, indicating undisturbed vertical exploratory behavior in TRα^GS^ mice (Fig. 3A). Latency to first entry into the center was higher in TRα^KO^ mice (Fig. 3B). Additionally, TRα^KO^ mice showed a decrease in the total number of entries into the center, center distance traveled, and time spent in the center, accompanied by an increase in time spent in the periphery (Fig. 3C-F). Interestingly, TRα^GS^ mice did not differ from their WT littermates in these anxiety-related indicators. Comparable results were obtained for female animals, although not every parameter reached statistical significance due to higher variation (Fig. S2).

**Figure 3:**
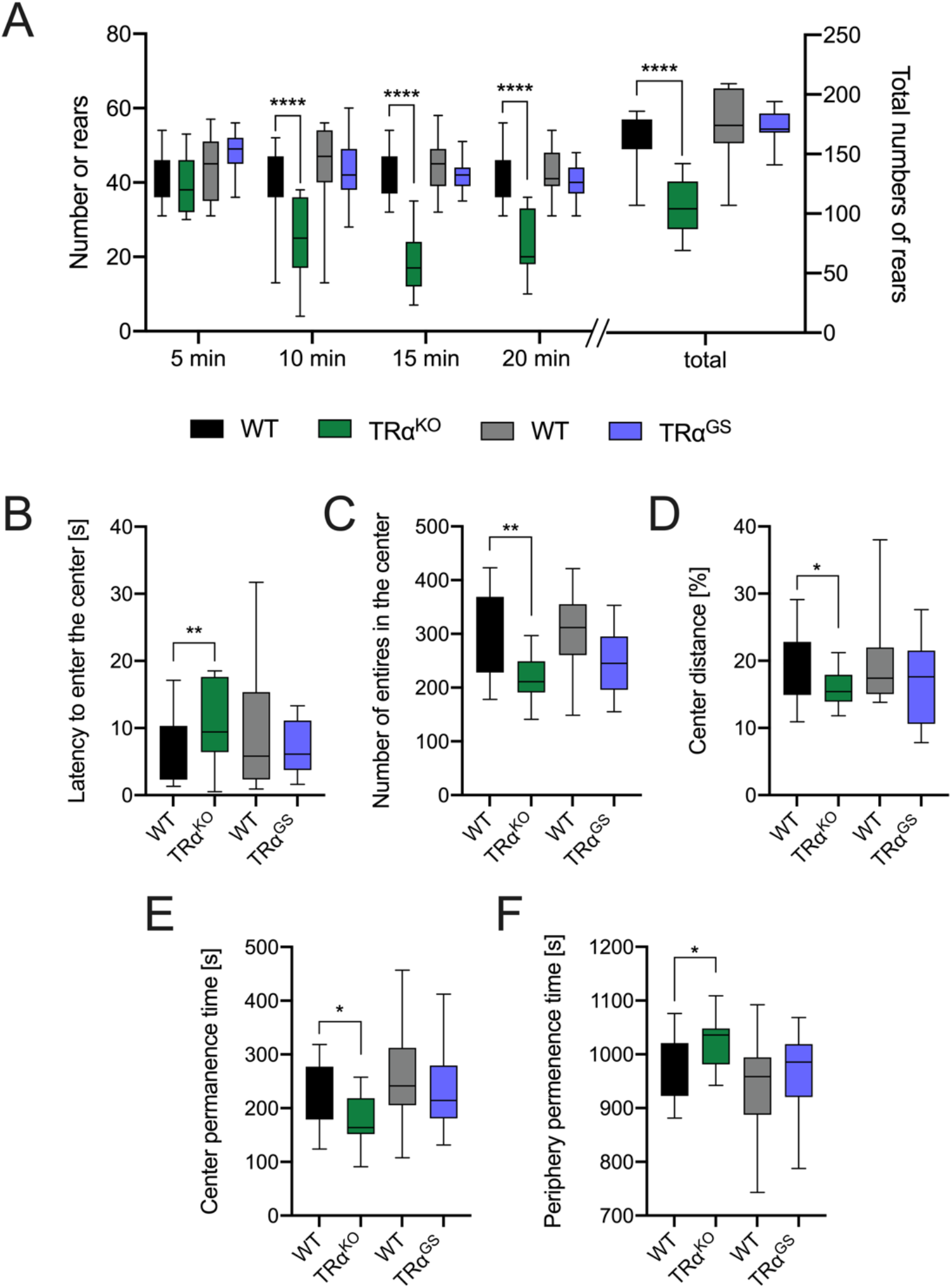
Open field test of male TRα mice. **A** Number of rears at certain time points and total number of rears (total) during 20 min test period. **B** Latency to enter the center. **C** Number of entries in the center. **D** Percentage of distance travelled in the center. **E** Time spent in the center and **F** time spent in the periphery. (One-way ANOVA with Sidak’s multiple comparison test; \**P<0.05*, \*\**P<0.01*, \*\*\**P<0.001*, \*\*\*\**P<0.0001*).

Previous studies have associated loss or impairment of TRα signaling with increased anxiety-related behavior in TRα^KO^ mice and mice with a dominant-negative, neuron-specific mutation of TRα ^33,34^. This behavior has been attributed to a reduced number of GABAergic interneurons and decreased density of GABAergic terminals in the CA1 region of the hippocampus ^35^. As TRα^GS^ mice did not display the hypoexploratory and anxiety-related features observed in TRα^KO^ mice, we hypothesize that central noncanonical TRα signaling plays a significant role in the regulation of behavior.

### Neurology and neuromuscular function

In the general observation series following a modified SHIRPA protocol, the only statistically significant difference observed was increased tail elevation in both sexes of TRα^KO^ mice compared to WT littermates (34.8% vs. 6.7%, respectively; P=0.014, Fisher’s Exact test) that was absent in the TRα^GS^ mice. However, no significant differences were observed in overall appearance, movement, and reflexes in both TRα and TRβ mutant mice compared to their WT littermates. Furthermore, measurement of grip strength, assessing muscle function, and evaluation of motor coordination and balance by Rotarod did not reveal any differences between TRα^KO^ and TRα^GS^ or TRβ^KO^ and TRβ^GS^ mice and their WT littermates.

The lack of a phenotype in muscle function and coordination in TRα^KO^ and TRα^GS^ mice was surprising as TRα is the predominant TR isoform in brain and muscle and hypothyroidism or defective TH transport into brain result in locomotor impairment ^36–38^. An explanation for the absence of a phenotype could be a TRβ-specific effect on locomotor activity. Mice that specifically express a dominant-negative TRβ (TRβG345R) in Purkinje cells had impaired cerebellar development resulting in significant disrupted motor coordination ^39^. Expression of a dominant-negative TRβ could also impair any TRα-mediated effects in these cells, thus this result does not allow determining the relevant TR isoform but rather helps to define the underlying cell-type. Global deletion of TRα1 prevented structural alterations of the Purkinje cell layer induced by hypothyroidism and these alterations can only be rescued by T3 and not GC-1, a TRβ specific agonist ^40^. This led to the conclusion that the impaired cerebellar development is rather the result of the apo-receptor function of TRα. This was further confirmed with TRαR384C mutant mice. The mutation results in a more than 10-fold lower affinity of TRα for T3 ^41^ and these mutant mice had locomotor dysfunctions that could be ameliorated by post-natal T3 injection, overcoming the reduced T3 affinity of the mutant receptor ^35^. Here, both the TRα^KO^ and the TRα^GS^ mice do not have any apo-receptor function as either the entire receptor is missing or the receptor lacks DNA-binding. Therefore, the absence of a neuromuscular phenotype in TRα^KO^ and TRα^GS^ mice can be explained by the lack of the apo-receptor function and thus rather reflects the phenotype of the TRα1^KO^ mice ^40^. This also implies that the role of TRα is redundant and can be compensated for grip strength and motor coordination, either by TRβ or other regulatory mechanisms.

### Metabolism

#### NMR measurements and indirect calorimetry

Body weight was slightly reduced in both TRα^GS^ and TRα^KO^ compared to WT mice with a more pronounced effect in males (Table 1). This weight reduction could primarily be attributed to decrease in fat mass accumulation in TRα^KO^ and TRα^GS^ mice compared to their WT littermates. Body composition analysis by two NMR measurements at 13 and 18 weeks confirmed that TRα^GS^ mice had less fat but preserved lean mass (Table 1). In TRα^KO^ mice both fat mass as well as lean mass were reduced, indicating that lean mass maintenance is linked to noncanonical TRα action. Interestingly, TRα^GS^ mice showed slightly but significantly decreased maximum oxygen consumption accompanied by an increased rearing behavior compared to their WT controls. However, in TRα^KO^ mice food intake and minimum oxygen consumption was increased, resulting in a higher respiratory exchange rate (RER)(Table 1). These findings collectively indicate that TRα mutants were in a mild hypermetabolic state, which was more pronounced in TRα^KO^ mice.

**Table 1:** NMR analysis and indirect calorimetry of male TRα mutant mouse lines.

| | WT | TR$\alpha^{KO}$ | | WT | TR$\alpha^{GS}$ | |
| --- | --- | --- | --- | --- | --- | --- |
| | mean ± SD | mean ± SD | p-value | mean ± SD | mean ± SD | p-value |
| avg. body mass [g] | 26.1 ± 1.8 | 22.3 ± 1.7 | < 0.001 | 25.8 ± 1.1 | 24.6 ± 1.4 | 0.003 |
| fat mass [g] | 5.5 ± 0.5 | 4.5 ± 0.6 | < 0.001 | 5.9 ± 0.5 | 4.6 ± 0.5 | < 0.001 |
| lean mass [g] | 17.9 ± 1.7 | 15.3 ± 1.2 | < 0.001 | 17.2 ± 1.0 | 17.3 ± 1.1 | 0.9690 |
| food intake [g] | 2.5 ± 1.0 | 3.0 ± 0.6 | 0.001 | 2.5 ± 0.8 | 2.6 ± 0.8 | 0.682 |
| avg. VO<sub>2</sub> [ml/(h*animal)] | 89.47 ± 11.19 | 83.49 ± 6.09 | 0.114 | 90.37 ± 4.00 | 86.89 ± 5.58 | 0.906 |
| min. VO<sub>2</sub> [ml/(h*animal)] | 60.93 ± 12.29 | 62.09 ± 6.76 | 0.003 | 66.07 ± 6.01 | 62.87 ± 6.35 | 0.796 |
| max. VO<sub>2</sub> [ml/(h*animal)] | 127.27 ± 15.87 | 109.46 ± 7.06 | 0.199 | 122.64 ± 5.53 | 114.73 ± 8.44 | 0.012 |
| avg. RER [VCO<sub>2</sub>/VO<sub>2</sub>] | 0.89 ± 0.05 | 0.93 ± 0.03 | < 0.001 | 0.88 ± 0.04 | 0.90 ± 0.04 | 0.488 |
| avg. distance [cm] | 3333 ± 1247 | 3179 ± 902 | 0.413 | 3358 ± 609 | 3191 ± 835 | 0.309 |
| avg. rearing [counts] | 88 ± 41 | 90 ± 48 | 0.15 | 114 ± 54 | 144 ± 77 | 0.017 |
| group size | n=15 | n=11 |  | n=14 | n=15 |  |

*Means ± SD and p-values of a Linear Model.*

These results are generally consistent with previous findings that TRα mutant mouse models showed hypermetabolism and defects in adipogenesis, resulting in resistance to obesity and a lean phenotype ^41–44^. However, there were differences: TRα^KO^ mice were hyperphagic, but TRα^GS^ mice were not. Furthermore, TRα^KO^ mice showed a shift towards carbohydrates as fuel for oxidative phosphorylation indicated by increased RER (Table 1). These data suggest that TRα^KO^ mice are much less efficient than TRα^GS^ mice in using energy sources, pointing to a role of noncanonical TRα signaling in energy metabolism.

We have previously reported that TRβ^GS^ mice exhibited increased locomotor activity and metabolic rate (higher minimal and average oxygen consumption) compared to TRβ^KO^ mice, whereas food consumption did not differ between genotypes, which we interpreted as a contribution of noncanonical TRβ signaling ^13^.

#### Glucose metabolism

In *ad libitum* fed animals, insulin levels were markedly decreased in male TRα^KO^ and TRα^GS^ mice (Fig. 4A). After a 6-hour food withdrawal, basal glucose levels were lower in TRα^GS^ mice compared to their respective controls (Fig. 4B), while no significant differences were observed for TRα^KO^ mice and their WT litter mates. However, an intraperitoneal glucose tolerance test (IPGTT) showed no differences in glucose tolerance between the mutants and the WT controls (Fig. 4C-E).

**Figure 4:**
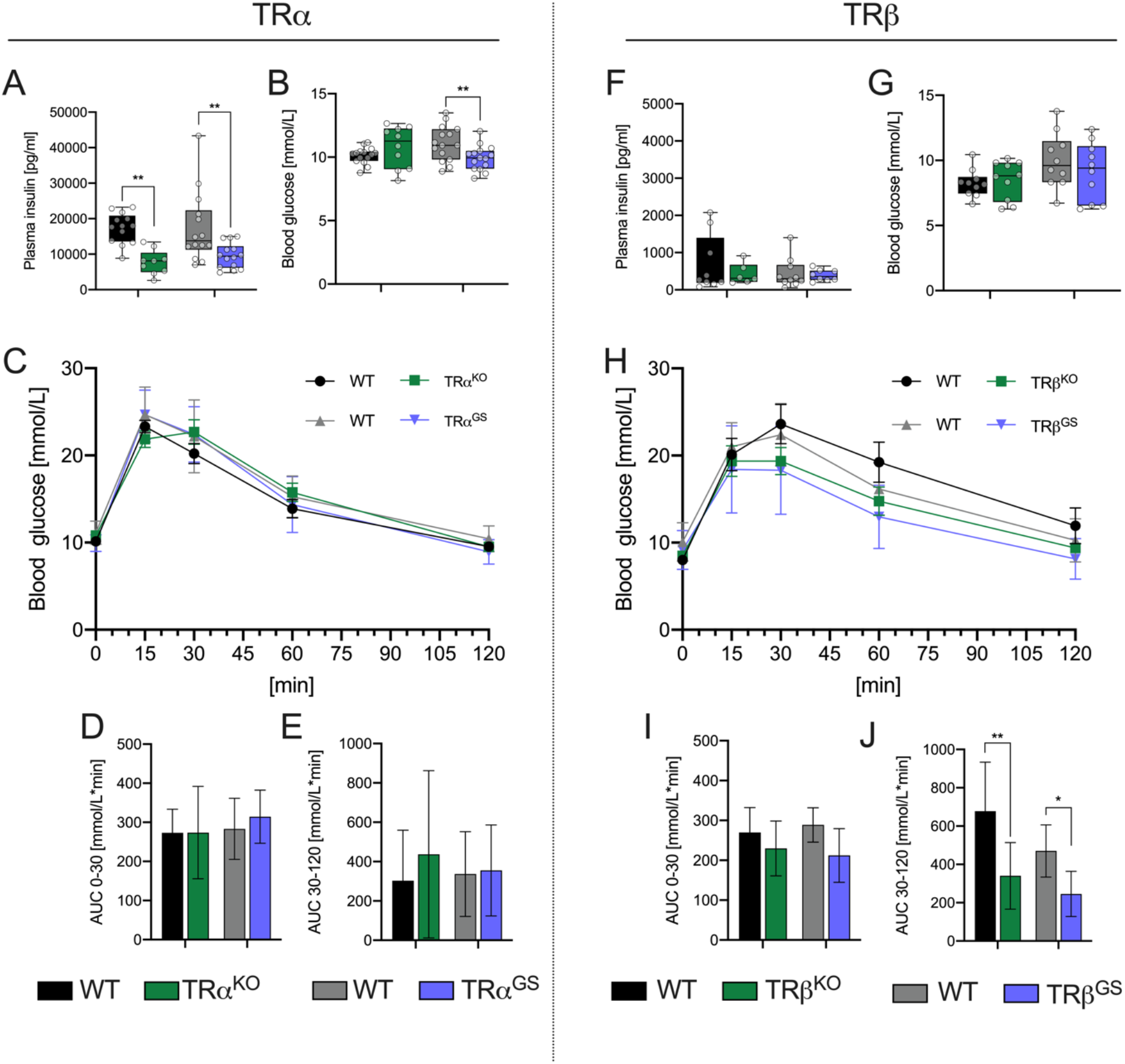
Intraperitoneal glucose tolerance test (IpGTT) in male and female TRα and TRβ mice. (A)Plasma insulin for male TRα (**A**) and male TRβ (**F**) mice. Blood glucose after 6h fasting at time point T0 of TRα (**B**) and TRβ (**G**) mice. IpGTT of TRα (**C**) and TRβ (**H**) mouse lines (mean±SEM). Area under the curve (AUC) of TRα mice for 0-30 min (D) and 30-120 min (E) and of TRβ mice (I-J). One-way ANOVA with Tukeýs *post-hoc* test; \**P<0.05*, \*\**P<0.01*; (n=11-15 TRα strains; n=8-10 TRβ strains).

In the TRβ strains, we found no major differences in basal insulin levels, determined in samples collected by final bleeding from *ad libitum* fed animals (Fig. 4F). However, both TRβ^KO^ and TRβ^GS^ mice exhibited improved glucose tolerance (lower area under the curve (AUC) glucose values during the 30-120 min interval, Fig. 4J).

TH regulates glucose homeostasis through actions in various organs such as pancreas, liver, gastrointestinal tract, adipose tissue, skeletal muscle, and the central nervous system ^45^. One explanation could be that canonical TRα action contributes to maintaining insulin concentration in the *ad libitum* fed condition. TH and TR signaling is important for pancreatic β-cell function and survival ^46,47^. Both main TRα and TRβ isoforms contribute to pancreatic homeostasis, with TRα stimulating β-cell proliferation and formation of insulin-producing cells ^48,49^, and TRβ improving survival and insulin secretion ^47,50,51^. A recent study highlighted the importance of TRα p43, a truncated isoform, in β-cell maturation and regulation of postnatal insulin content ^52^. Therefore, the loss of canonical TRα action by the GS-mutation, which also affects the truncated p43 isoform, may lead to a decreased β-cell number and subsequently in reduction in serum insulin concentrations. However, a decreased β-cell number is not well compatible with preserved glucose tolerance and unaltered or even lower glucose concentrations during fasting in these animals. Therefore, a better explanation would be an increased glucose metabolism due to an increased overall energy demand in TRα^KO^ and TRα^GS^ mice. TRα mutant mice, KO as well as GS, have lower body weights, especially fat mass, due to hypermetabolism, which supports this hypothesis. Therefore, the lower insulin concentrations possibly reflect improved glucose tolerance and leanness.

The loss of canonical TRβ action in both TRβ^KO^ or TRβ^GS^ mice resulted in improved glucose tolerance. Both TRβ mutants are resistant to TH due to impaired negative feedback on the HPT axis, resulting in elevated TH serum concentrations ^12,13^. Therefore, the improved glucose tolerance in both TRβ mutant mouse lines can probably be attributed to increased TH effects on tissues predominantly expressing TRα, including adipocytes ^53,54^, skeletal muscle ^55,56^ and other cell types like thymocytes ^57^, by promoting the expression of glucose transporters and their translocation to the membrane. The pancreatic β-cells seem not to be affected, as insulin concentrations were not changed in these animals. Of note, acute glucose uptake after T3 treatment requires noncanonical TRβ signaling ^13^.

#### Lipid (triglyceride) metabolism

In our previous study, we reported elevated serum triglyceride (TG) concentration in TRβ^KO^ mice, while TG concentration in TRβ^GS^ mice were comparable to those of WT mice ^13^. In the current screening, we observed trends towards the expected direction in TG concentrations for both TRβ^KO^ and TRβ^GS^ mutants, but they did not reach statistical significance (Fig. S3). Fasting TG concentrations were reduced in both TRα^KO^ and TRα^GS^ mutants compared to their respective WT controls following overnight fasting (Table 2), possibly again a result of inefficient metabolism. Non-esterified fatty acids (NEFA) and glycerol levels were not affected. We observed a genotype-specific difference in fasting cholesterol, with slightly increased levels in TRα^GS^ mutants due to elevated non-HDL cholesterol, while fasting cholesterol was decreased in TRα^KO^ mice primarily due to lower HDL-cholesterol levels. These effects were modest.

**Table 2:** Serum fatty acid parameters under fasting conditions.

| | WT | TRα<sup>KO</sup> | | WT | TRα<sup>GS</sup> | |
| --- | --- | --- | --- | --- | --- | --- |
| | mean ± SD | mean ± SD | p-value | mean ± SD | mean ± SD | p-value |
| fasting Glucose [mmol/l] | 6.57 ± 0.86 | 8.92 ± 1.26 | < 0.001 | 7.17 ± 1.07 | 8.25 ± 2.08 | 0.191 |
| fasting Cholesterol [mmol/l] | 2.432 ± 0.234 | 2.234 ± 0.412 | 0.06 | 2.521 ± 0.328 | 2.603 ± 0.209 | 0.049 |
| fasting HDL-cholesterol [mmol/l] | 1.908 ± 0.162 | 1.684 ± 0.318 | 0.005 | 1.976 ± 0.209 | 2.036 ± 0.177 | 0.091 |
| fasting non-HDL Cholesterol | 0.524 ± 0.083 | 0.55 ± 0.139 | 0.35 | 0.545 ± 0.128 | 0.567 ± 0.063 | 0.035 |
| fasting Triglycerides [mmol/l] | 0.825 ± 0.189 | 0.558 ± 0.151 | < 0.001 | 0.88 ± 0.233 | 0.732 ± 0.184 | 0.017 |
| fasting NEFA [mmol/l] | 1.15 ± 0.12 | 1.02 ± 0.28 | 0.14 | 1.18 ± 0.23 | 1.23 ± 0.29 | 0.972 |
| fasting Glycerol [mmol/l] | 0.247 ± 0.03 | 0.278 ± 0.091 | 0.76 | 0.244 ± 0.048 | 0.271 ± 0.072 | 0.433 |
| group size | n=15 | n=11 |  | n=14 | n=15 |  |

*Means ± SD and p-values of a Linear Model.*

Some previously observed phenotypes, such as elevated TG levels in TRβ^KO^ mice ^13^, were not confirmed as statistically significant in the present data. This could be attributed to a phenotypic heterogeneity due to the mixed 129-C57BL/6 background, as these two strains differ significantly in terms of energy metabolism ^58^. Although efforts were made to minimize background effects through eight generations of backcrossing to a pure background, residual background effects may still exist if relevant genes are closely genetically linked to the site of the investigated mutation.

### Clinical Chemistry

*Ad libitum* fed plasma proteins, enzyme activities, and electrolytes were assessed. The majority of observed genotype effects were moderate and consistent in both TRα mutant lines, indicating their mediation through the canonical pathway. For example, in female and male TRα mutant mice, alkaline phosphatase (ALP) activity and lactate levels were significantly increased (Fig. 5A & B). Leptin levels were decreased in both mutant lines (Fig. 5C). There were a few exceptions to these trends. Specifically, plasma albumin concentration was mildly decreased in TRα^GS^ mice but unchanged in TRα^KO^ mice. (Fig. 5D). Creatinine levels were increased in TRα mutants, with a more pronounced effect in TRα^GS^ mice (Fig. 5E). Interestingly, the opposite trend was observed in TRβ^GS^ mice, where creatinine levels were decreased (Fig. 5F). However, no significant effects on other clinical chemistry parameters were observed in TRβ mutant animals. Iron levels showed only minor changes in TRα^KO^ mutant mice, with no significant alterations in unsaturated iron binding capacity (UIBC). However, total iron binding capacity (TIBC) was noticeably increased in TRα^KO^ mutant mice (Fig. 5G). In contrast, neither iron levels nor UIBC and TIBC were significantly affected in TRα^GS^ mice. The results from clinical chemistry parameters in mice with TR mutations provide insights into the physiological effects of the TRs and their signaling. The increase in lactate concentration in mice lacking the canonical TRα signaling pathway suggests a shift towards anaerobic muscle metabolism. In rats, T3 treatment enhanced glycolysis, influenced mitochondrial uncoupling, and affected mitochondrial substrate utilization in skeletal muscle ^59^. T3-induced increase in energy expenditure in skeletal muscle required TRα in mice ^25^. The elevated lactate concentrations observed in TRα mutant mice may reflect alterations in muscle metabolism and energy utilization, e.g. increased glycolysis.

**Figure 5:**
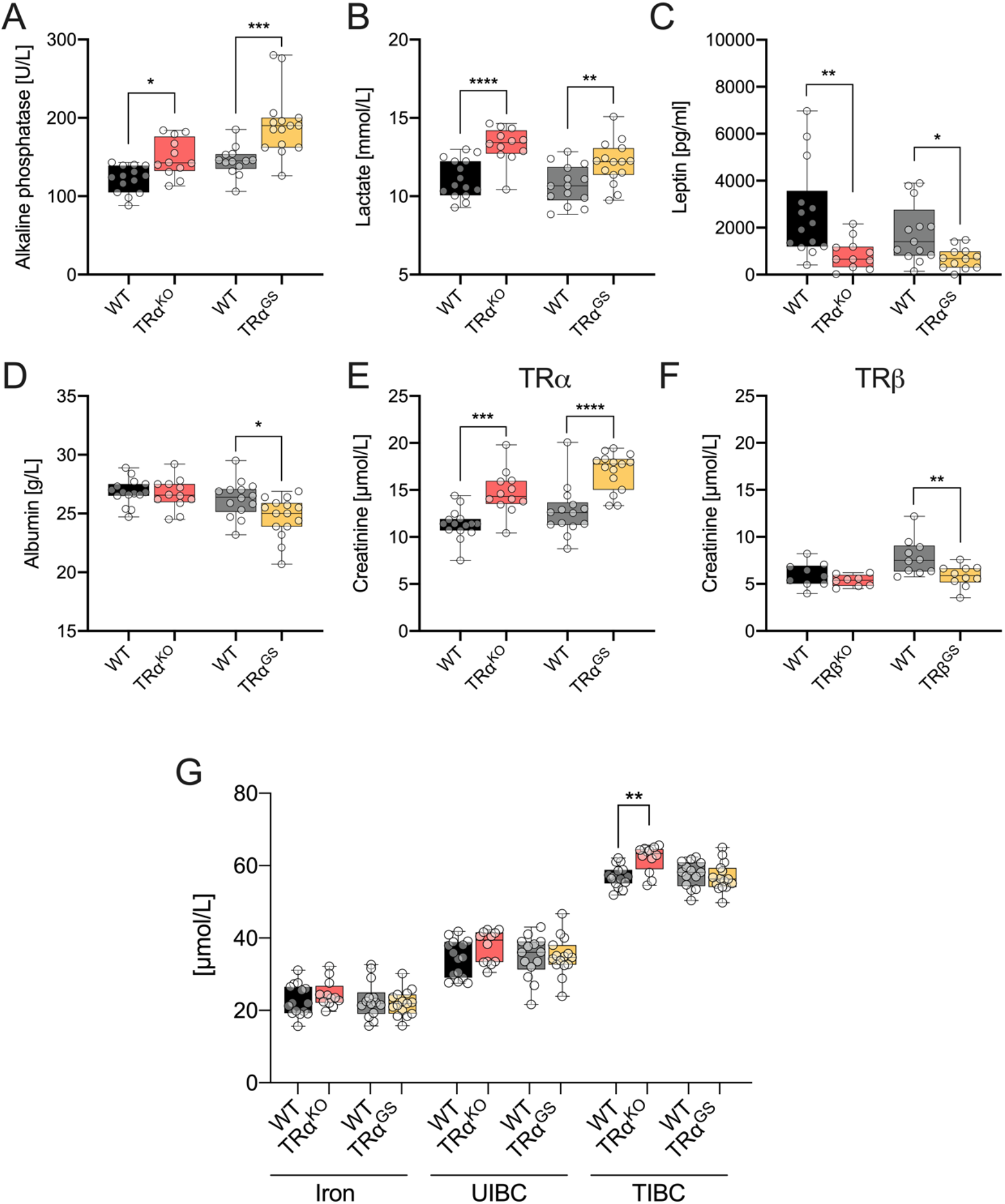
Clinical chemistry –Plasma proteins, enzyme activities, and electrolytes (*ad libitum* fed mice). For female TRα mouse lines plasma activity of alkaline phosphatase **A**, concentration of lactate **B**, leptin **C**, and albumin **D** are shown. Creatinine levels are depicted for TRα **E** and TRβ **F** mice. **G** Iron concentration, unsaturated iron binding capacity (UIBC) and total iron binding (capacity TIBC) of female TRα mouse lines. One-way ANOVA; \**P<0.05*, \*\**P<0.01*, \*\*\**P<0.001, ****P<0.0001*; (n=11-15 TRα strains; n=8-10 TRβ strains).

The elevated serum creatinine levels in TRα mutants also indicate metabolic changes in skeletal muscle, which experience sudden demands for energy ^60^. However, it is important to note that the accumulation of plasma creatinine could also be attributed to impaired renal clearance, as thyroid hormone status affects renal function ^61,62^.

The lower concentrations of leptin in the plasma of TRα^KO^ and TRα^GS^ mutants can be attributed to the reduced white adipose tissue (WAT) mass observed in these animals, as plasma leptin levels are known to correlate with adipose mass ^63^. However, the relationship between thyroid hormone and leptin is complex. Some studies reported elevated leptin levels in hypothyroidism and reduced levels in hyperthyroidism, correlating with body mass index (BMI) and thyroid-stimulating hormone (TSH) levels ^64^. In contrast, another study found no correlation between leptin levels, BMI, or TSH, but instead correlated leptin with visceral adipose tissue mass ^65^. Thus, the specific mechanisms linking TH/TR signaling and leptin regulation require further investigation.

Only TRα^GS^ mice, male and female, had reduced albumin serum concentrations. While we cannot with certainty explain this finding, a smaller liver size may contribute to this effect. Liver weight as well as the liver to body weight and the liver to tibia length ratio were significantly decreased in female (*P*=0.016) and male (*P*=0.032) TRα^GS^ mice. Interestingly, no such changes were observed for TRα^KO^ mice and their albumin concentration was the same as that in WT mice.

Regarding iron metabolism, in cases of iron-deficiency anemia, total iron binding capacity (TIBC) is increased. Similarly, TRα^KO^ mice with mild macrocytic anemia (see below) had an increased TIBC, which suggests a compensatory mechanism ^66^.

### Hematology and immunology

#### Hematology

Red blood count parameters in TRβ^KO^ and TRβ^GS^ mice were similar to WT (Fig S4). In TRα^KO^ mice red blood cell count was reduced (Fig. 6A) while mean corpuscular volume (MCV) and mean corpuscular hemoglobin content (MCH) were increased (Fig. 6B & C). Thus, the TRα^KO^ phenotype resembled macrocytic anemia. In contrast, these parameters were not altered in TRα^GS^ mice compared to their WT littermates (Fig. 6A-C). These data suggest that the presence of noncanonical TRα action suffices to support normal erythrocyte development and maturation.

**Figure 6:**
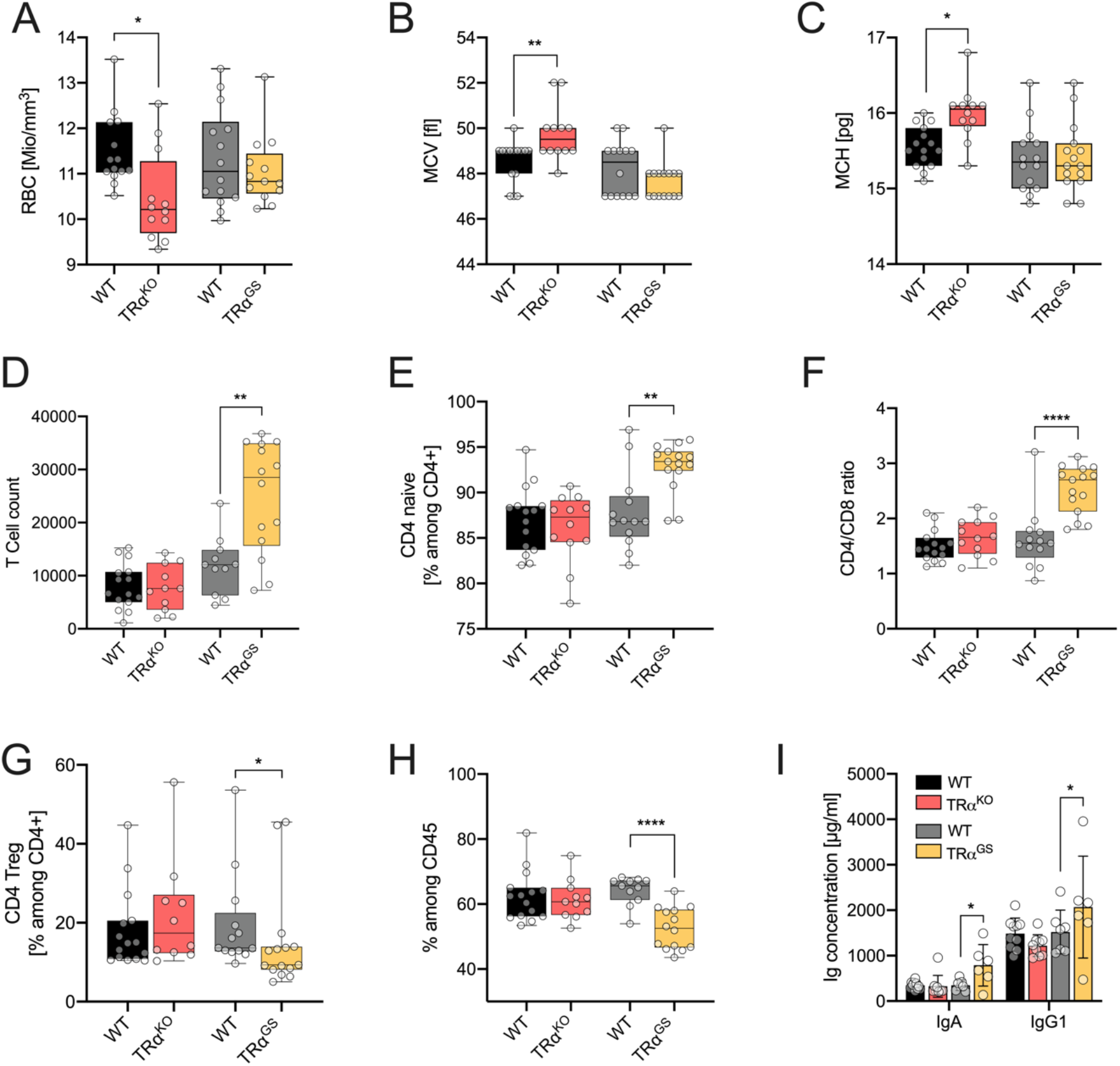
Hematological and immunological analysis of female TRα mice. For female TRα mouse lines red blood cell count (RBC) **A**, mean corpuscular volume (MCV) **B** and the mean corpuscular hemoglobin content (MCH) **C** were determined. **D** Total number of T cells. **E** Percentage of the CD4+,CD62l+,CD44-, T cell subpopulation. **F** Ratio CD4/CD8 T cells. **G** Percentage of the CD4+,CD25+, T cell subpopulation. **H** Percentage of CD45+ B cells. **I** Plasma concentration of immunoglobulins IgA and IgG1. A, B, C and I: One-way ANOVA; D, E, F, G, H, Mann Whitney test; \**P<0.05*, \*\**P<0.01*, \*\*\**P<0.001, ****P<0.0001*; (n=11-15).

The stimulatory influence of TH on erythropoiesis had long been recognized in epidemiological studies. Hemoglobin concentration was lower in hypothyroid patients than in euthyroid participants and TH concentration even predicted future anemia ^67,68^. Significant positive linear relationships existed between free TH and hemoglobin, hematocrit and erythrocyte count ^69^. Strikingly, lower T3 concentration was associated with higher MCV and MCH, matching the phenotype in TRα^KO^ mice.

Mouse studies could attribute the TH effect on erythropoiesis to TRα. Adult TRα^KO^ mice had lower hematocrit than WT mice and T3 and TRα stimulated development and maturation of spleen erythroblasts ^70,71^. Spleens from TRα^KO^ mice contained more immature erythroblasts, indicating impaired terminal differentiation. A mouse model of RTHα, the TRα^PV/+^ mouse, showed reduced RBC, hemoglobin and hematocrit ^72^. Terminal differentiation of TRα^PV/+^ of bone marrow cells into erythrocytes, especially enucleation, was impaired. The relevance of TRα was also found in humans because most patients with RTHα show mild anemia ^73,74^. PI3K activation is the best characterized mode of noncanonical TRα action ^6^. In this context it is interesting that the PI3K signaling pathway also is an important mediator of erythrocyte proliferation and terminal differentiation including polarization of erythroblasts and enucleation ^75^. Our data indicate that noncanonical TRα action, possibly via PI3K activation, mediates the influence of TH on erythropoiesis and erythrocyte differentiation.

#### Immunology

The immunological phenotype of TRα^KO^, TRα^GS^, TRβ^KO^ and TRβ^GS^ mice was examined using flow cytometry to determine the frequencies of leukocyte populations in peripheral blood. Interestingly, elevated total numbers of T cells were found in female and male TRα^GS^, while the T lymphocyte populations of TRα^KO^ mice showed no significant alterations (Fig. 6D). The increase in T lymphocytes is predominantly caused by an elevation in CD4+ T cells, the majority of which are expressing CD62L+ CD44-, characterizing them as naive cells (Fig. 6E,F), F & G). Similarly, a reduction in CD4+ CD25+ regulatory T cells has been observed (Fig. 6G). However, the analysis of these animals has been in unchallenged conditions, and although it may not necessarily imply a pathological state, it could be crucial in an infection or challenge of the immune system. Despite reduced frequencies of B cell subpopulations in TRα^GS^ mice, with only percentages changed potentially due to the increase in T cells (Fig. 6H), we found higher concentrations of immunoglobulins (Ig), particularly IgA and IgG1, which are associated with gastrointestinal inflammatory diseases or autoimmune conditions (Fig. 6I).

The effect of TH on the immune system has already been the subject of previous studies ^76–78^. However, the results obtained so far are conflicting, showing both inhibitory and stimulatory actions of TH within the same type of immune cells. Furthermore, the underlying mechanisms, including the contribution of TRα and TRβ signaling, which are expressed in immune cells, remain elusive. Regarding T cell immunity, previous studies have demonstrated reduced T cell numbers in TRα^KO^ mice ^79^. Conversely, enhanced T cell proliferation and anti-tumor immunity were observed after T4 treatment in a murine lymphoma model ^80,81^. On the contrary, TH were induced T cell apoptosis upon *in vitro* stimulation of human peripheral blood lymphocytes ^82^. Importantly, previous studies did not distinguish between different T cell subtypes, such as CD4+ and CD8+ T cells, and therefore may have reported conflicting results. Here, we report a differential action of TH in CD4+ and CD8+ T cell subsets, as well as a potential prominent role of noncanonical TRα action in CD4+ but not CD8+ T cells. In addition to T lymphocytes, previous studies have found reduced frequencies of B cells in peripheral blood of naïve TRα^KO^ mice and a murine model of TRα resistance (TRα1^PV/+^), indicating an essential role of TRα for primary B cell development ^79,83^. Consistent with these findings, our results from TRα^GS^ mice suggest that canonical TRα action is relevant for lymphocyte development. Additionally, it should be noted that the elevated TH serum levels in TRβ mutant mice may lead to a desensitizing effect on TRα-mediated signaling, potentially resulting in a phenotype similar to that of TRα mutant mice ^84,85^. The present data suggest a differential role of noncanonical and canonical TRα signaling in CD4+ and CD8+ T cells, while primary B cell development appears to rely on canonical TRα actions. However, further studies, ideally with T3 treatment and infection models, are necessary to precisely define the impact of noncanonical and canonical TRα signaling on T and B cell immunity.

## Conclusion

Aims of this genotype comparison were to determine for which physiological effects TR action is relevant and to attribute these effects to the TRα or TRβ isoform and their canonical or noncanonical mode of action. Pronounced perturbations driven by the absence of canonical TR action were mostly found for the senses, especially hearing (mainly TRβ, to a lesser degree TRα), visual acuity and retinal thickness (TRα and TRβ), for muscle metabolism (TRα) and heart rate (TRα).

Interestingly, for several physiological effects, selective abrogation of canonical TR action did not produce a phenotype reminiscent of the full KO. Unlike TRα^KO^ mice, TRα^GS^ mice did not show anemia, reduced retinal vascularization or increased anxiety related behavior, indicating that noncanonical TR action sufficed to maintain the WT phenotype for these effects. Furthermore, TRα^KO^ mice were leaner than WT mice despite hyperphagia, demonstrating the resistance to obesity and inefficient utilization of energy from food ^42^. TRα^GS^ mice, in contrast to TRα^KO^ mice, did not exhibit the same degree of leanness and did not consume more food than their respective WT controls. The preservation of their noncanonical TRα action suggests more efficient energy utilization. However, the specific organ or cell type, as well as the biochemical pathway through which noncanonical TRα action influences metabolic efficiency, remains to be determined. From these findings, we may conclude that noncanonical TR action contributes to the diverse spectrum of TR effects in physiology.

More mechanistic conclusions can be drawn from a comparison of present data with the phenotype of RTHα patients that show mild anemia ^73^. Similarly, TRα^KO^ mice, but not TRα^GS^ mice, showed mild anemia, indicating that noncanonical TRα action sufficed to produce the WT phenotype and contribute to normal erythrocyte maturation. We hypothesize that the preserved PI3K activation, which is one mode of noncanonical TRα action, mediates terminal differentiation of erythroblasts and enucleation ^75^. Similarly, hepatic TG content is regulated by TRβ, increased in T3-resistant TRβPV mice and RTHβ patients, but not in mice with preserved noncanonical TRβ action (TRβ^GS^) ^13,86,87^. From these results, we conclude that the dominant negative effect of TRα and TRβ is not restricted to their canonical action and gene expression, but also extends to their noncanonical action. The current concept on noncanonical TR activation of the PI3K pathway assumes that the TRs bind the regulatory subunit of PI3K, p85α, and release it after T3 binding ^7,88^, which leads to AKT phosphorylation and activation of the pathway. Impaired T3 binding thus would leave p85α bound to the unliganded TR, effectively inactivating p85α.

It is tempting to speculate that the phenotype of potential human TR mutations that exclusively abrogate canonical signaling would differ from classical RTH with mutation that impair hormone binding due to preserved noncanonical TR signaling. Based on the present data, we hypothesize that mutations abrogating only DNA-binding of TRα, a ‘canonical-only RTHα’, would not result in anemia and hypermetabolism, and anxiety would be less severe compared to classical RTHα.

In general, the mice with absent canonical (GS) or both canonical and noncanonical TR action (KO) presented a rather mild phenotype. This may be explained by several mechanisms. For example, absence of one TR isoform could be compensated by the other isoform, which would prevent an overt phenotype. Furthermore, the relative lack of effects in TR KO mice does not necessarily indicate that TH/TRs are not involved in regulating key physiological processes. Rather, it indicates that those processes are redundantly regulated starting at germline development, and absence of TR is compensated by other hormones, receptors, and pathways. Studies of mice with a dominant negative TR mutant, such as TRαAMI, TRαPV or TRβPV mice, would address this and reveal TR involvement. However, dominant-negative effects of one isoform could impair effects where TRs play only a minor role and could spill over to functions of the other isoform, whereas our approach reveals physiological effects where TRs are crucially involved. Thus, when studying TR KO and dominant-negative TR mutant mice, the possibility of under- and overestimation of TR effects needs to be considered. Another reason for the mild phenotype could be that we studied untreated mice. Some physiologically significant changes or their absence may become apparent only after T3 treatment.

Here, we studied global KO and knock-in (KI) mice. A future challenge is to determine the cell-type where TH/TRs act, possibly in cell-type specific KO and KI mice, both untreated and with T3 treatment in comparison. One aspect to consider in interpreting the results here is that they may include effects of altered TR action during development. Overcoming this requires inducible TR KO and KI mouse models. Cell type-specific models would also help to distinguish direct from remote TR effects, which occur on peripheral organs after central TH/TR action ^43,89,90^. Clearly, experiments addressing cell-type specificity, developmental aspects and T3-treatment will be reserved for focused questions that could derive from the present data, e.g. studying erythrocyte maturation or energy metabolism.

In summary, the TRα and TRβ KO and GS comparison demonstrates that canonical and noncanonical actions of TRα and TRβ contribute to various physiological processes, e.g. retinal development, anxiety-related behavior, body composition, metabolic efficiency, muscle metabolism, heart rate, hematological parameters, and immune cell populations. These data underscore the importance of both canonical and noncanonical TR signaling pathways in different aspects of physiology and the need for more detailed investigations of the precise mechanisms for each effect.

## Acknowledgements

We thank Prof. Dr. G. Hilken, Dr. A. Wißmann, Dr. P. Dammann and the staff of the core animal facility at the University Hospital Essen for their continued dedicated support. We acknowledge the expert technical support of our technicians and of the Core Facility Laboratory Animal Services at Helmholtz Munich.

## Funding

This work was supported by Deutsche Forschungsgemeinschaft (DFG, German Research Foundation) grant MO1018/2-2 (L.C.M.) and Project-ID 424957847 TRR 296 LOCOTACT (P.T.P., T.D.M., D.F. & L.C.M.), and by an IFORES grant from the Faculty of Medicine, University of Duisburg-Essen and by the German Federal Ministry of Education and Research (Infrafrontier grant 01KX1012 to M.H.A.) and the German Center for Diabetes Research (DZD) (M.H.A.).

## Author Contributions

L.C.M. conceived the study. M.H.A., V.G.D. and H.F. conceptualized the phenotyping tests. G.S.H., D.G., C.W., L.B., O.V.A., L.G., B.R., J.R., N.S. and I.T. conducted animal breeding, genotyping, experiments and acquisition of data. V.G.D., H.F., W.W., E.W., D.F. and L.C.M. supervised the study, V.G.D. and H.F. managed and coordinated the project at the GMC. A.A.P., N.D., L.B., O.V.A., L.G.. S.M.H., B.R., J.R., N.S., I.T., P.T.P., T.D.M., G.S.H., D.G., C.W., D.F. and L.C.M. analyzed and interpreted the data. G.S.H., D.G., C.W. and L.C.M. wrote the manuscript and all authors contributed to the final version. Funding and resources for the study were acquired by M.H.A., P.T.P., T.D.M., D.F. and L.C.M.

## Notes

### Competing Interest Statement

The authors have declared no competing interest.

## References

1. Yen PM. Physiological and molecular basis of thyroid hormone action. Review. Physiological reviews. Jul 2001;81(3):1097–142. doi:10.1152/physrev.2001.81.3.1097

2. Flamant F, Cheng SY, Hollenberg AN, et al. Thyroid Hormone Signaling Pathways: Time for a More Precise Nomenclature. Endocrinology. Jul 1 2017;158(7):2052–2057. doi:10.1210/en.2017-00250

3. Singh BK, Sinha RA, Yen PM. Novel Transcriptional Mechanisms for Regulating Metabolism by Thyroid Hormone. International journal of molecular sciences. Oct 22 2018;19(10)doi:10.3390/ijms19103284

4. Cao X, Kambe F, Moeller LC, Refetoff S, Seo H. Thyroid hormone induces rapid activation of Akt/protein kinase B-mammalian target of rapamycin-p70S6K cascade through phosphatidylinositol 3-kinase in human fibroblasts. Mol Endocrinol. Jan 2005;19(1):102–12. doi:10.1210/me.2004-0093

5. Cao X, Kambe F, Yamauchi M, Seo H. Thyroid-hormone-dependent activation of the phosphoinositide 3-kinase/Akt cascade requires Src and enhances neuronal survival. The Biochemical journal. Dec 1 2009;424(2):201–9. doi:10.1042/BJ20090643

6. Hiroi Y, Kim HH, Ying H, et al. Rapid nongenomic actions of thyroid hormone. Research Support, N.I.H., Extramural Research Support, Non-U.S. Gov’t. Proceedings of the National Academy of Sciences of the United States of America. Sep 19 2006;103(38):14104–9. doi:10.1073/pnas.0601600103

7. Martin NP, Marron Fernandez de Velasco E, Mizuno F, et al. A rapid cytoplasmic mechanism for PI3 kinase regulation by the nuclear thyroid hormone receptor, TRbeta, and genetic evidence for its role in the maturation of mouse hippocampal synapses in vivo. Endocrinology. Sep 2014;155(9):3713–24. doi:10.1210/en.2013-2058

8. Kalyanaraman H, Schwappacher R, Joshua J, et al. Nongenomic thyroid hormone signaling occurs through a plasma membrane-localized receptor. Science signaling. May 20 2014;7(326):ra48. doi:10.1126/scisignal.2004911

9. Gauthier K, Plateroti M, Harvey CB, et al. Genetic analysis reveals different functions for the products of the thyroid hormone receptor alpha locus. Research Support, Non-U.S. Gov’t Research Support, U.S. Gov’t, P.H.S. Mol Cell Biol. Jul 2001;21(14):4748–60. doi:10.1128/MCB.21.14.4748-4760.2001

10. Brent GA. Tissue-specific actions of thyroid hormone: insights from animal models. Reviews in endocrine & metabolic disorders. Jan 2000;1(1-2):27–33.

11. Shibusawa N, Hollenberg AN, Wondisford FE. Thyroid hormone receptor DNA binding is required for both positive and negative gene regulation. J Biol Chem. Jan 10 2003;278(2):732–8. doi:10.1074/jbc.M207264200

12. Shibusawa N, Hashimoto K, Nikrodhanond AA, et al. Thyroid hormone action in the absence of thyroid hormone receptor DNA-binding in vivo. Journal of Clinical Investigation. 2003;112(4):588–597. doi:10.1172/jci200318377

13. Hones GS, Rakov H, Logan J, et al. Noncanonical thyroid hormone signaling mediates cardiometabolic effects in vivo. Proceedings of the National Academy of Sciences of the United States of America. Dec 26 2017;114(52):E11323–E11332. doi:10.1073/pnas.1706801115

14. Geist D, Hones GS, Gassen J, et al. Noncanonical Thyroid Hormone Receptor alpha Action Mediates Arterial Vasodilation. Endocrinology. Jul 1 2021;162(7)doi:10.1210/endocr/bqab099

15. Gauthier K, Chassande O, Plateroti M, et al. Different functions for the thyroid hormone receptors TRalpha and TRbeta in the control of thyroid hormone production and post-natal development. Research Support, Non-U.S. Gov’t. The EMBO journal. Feb 1 1999;18(3):623–31. doi:10.1093/emboj/18.3.623

16. Gailus-Durner V, Fuchs H, Becker L, et al. Introducing the German Mouse Clinic: open access platform for standardized phenotyping. Nature methods. Jun 2005;2(6):403–4. doi:10.1038/nmeth0605-403

17. Fuchs H, Aguilar-Pimentel JA, Amarie OV, et al. Understanding gene functions and disease mechanisms: Phenotyping pipelines in the German Mouse Clinic. Behavioural brain research. Oct 15 2018;352:187–196. doi:10.1016/j.bbr.2017.09.048

18. Fuchs H, Gailus-Durner V, Adler T, et al. Mouse phenotyping. Methods (San Diego, Calif). Feb 2011;53(2):120-35. doi:10.1016/j.ymeth.2010.08.006

19. Rathkolb B, Fuchs H, Gailus-Durner V, Aigner B, Wolf E, Hrabě de Angelis M. Blood Collection from Mice and Hematological Analyses on Mouse Blood. Curr Protoc Mouse Biol. Jun 1 2013;3(2):101–19. doi:10.1002/9780470942390.mo130054

20. Rathkolb B, Hans W, Prehn C, et al. Clinical Chemistry and Other Laboratory Tests on Mouse Plasma or Serum. Curr Protoc Mouse Biol. Jun 1 2013;3(2):69–100. doi:10.1002/9780470942390.mo130043

21. Moreth K, Fischer R, Fuchs H, et al. High-throughput phenotypic assessment of cardiac physiology in four commonly used inbred mouse strains. Journal of comparative physiology B, Biochemical, systemic, and environmental physiology. Aug 2014;184(6):763–75. doi:10.1007/s00360-014-0830-3

22. Pawliczek D, Dalke C, Fuchs H, et al. Spectral domain -Optical coherence tomography (SD-OCT) as a monitoring tool for alterations in mouse lenses. Experimental eye research. Jan 2020;190:107871. doi:10.1016/j.exer.2019.107871

23. Rozman J, Rathkolb B, Neschen S, et al. Glucose tolerance tests for systematic screening of glucose homeostasis in mice. Curr Protoc Mouse Biol. Mar 2 2015;5(1):65–84. doi:10.1002/9780470942390.mo140111

24. Forrest D, Erway LC, Ng L, Altschuler R, Curran T. Thyroid hormone receptor beta is essential for development of auditory function. Research Support, Non-U.S. Gov’t Research Support, U.S. Gov’t, P.H.S. Nat Genet. Jul 1996;13(3):354–7. doi:10.1038/ng0796-354

25. Nicolaisen TS, Klein AB, Dmytriyeva O, et al. Thyroid hormone receptor α in skeletal muscle is essential for T3-mediated increase in energy expenditure. The FASEB Journal. Sep 23 2020;.,,,,doi:10.1096/fj.202001258RR

26. Harvey CB, O’Shea PJ, Scott AJ, et al. Molecular mechanisms of thyroid hormone effects on bone growth and function. Research Support, Non-U.S. Gov’t Review. Mol Genet Metab. Jan 2002;75(1):17–30. doi:10.1006/mgme.2001.3268

27. Affortit C, Blanc F, Nasr J, et al. A disease-associated mutation in thyroid hormone receptor α1 causes hearing loss and sensory hair cell patterning defects in mice. Science signaling. Jun 14 2022;15(738):eabj4583. doi:10.1126/scisignal.abj4583

28. Winter H, Ruttiger L, Muller M, et al. Deafness in TRbeta mutants is caused by malformation of the tectorial membrane. The Journal of neuroscience : the official journal of the Society for Neuroscience. Feb 25 2009;29(8):2581–7. doi:10.1523/JNEUROSCI.3557-08.2009

29. Yang F, Ma H, Ding XQ. Thyroid Hormone Signaling in Retinal Development, Survival, and Disease. Vitamins and hormones. 2018;106:333–349. doi:10.1016/bs.vh.2017.05.001

30. Aramaki M, Wu X, Liu H, et al. Transcriptional control of cone photoreceptor diversity by a thyroid hormone receptor. Proceedings of the National Academy of Sciences of the United States of America. Dec 6 2022;119(49):e2209884119. doi:10.1073/pnas.2209884119

31. Sevilla-Romero E, Muñoz A, Pinazo-Durán MD. Low thyroid hormone levels impair the perinatal development of the rat retina. Ophthalmic Res. Jul-Aug 2002;34(4):181–91. doi:10.1159/000063885

32. Sjöberg M, Vennström B, Forrest D. Thyroid hormone receptors in chick retinal development: differential expression of mRNAs for alpha and N-terminal variant beta receptors. Development. Jan 1992;114(1):39–47. doi:10.1242/dev.114.1.39

33. Richard S, Aguilera N, Thevenet M, Dkhissi-Benyahya O, Flamant F. Neuronal expression of a thyroid hormone receptor alpha mutation alters mouse behaviour. Behavioural brain research. Dec 20 2016;doi:10.1016/j.bbr.2016.12.025

34. Wilcoxon JS, Nadolski GJ, Samarut J, Chassande O, Redei EE. Behavioral inhibition and impaired spatial learning and memory in hypothyroid mice lacking thyroid hormone receptor alpha. Behavioural brain research. Feb 12 2007;177(1):109–16. doi:10.1016/j.bbr.2006.10.030

35. Venero C, Guadano-Ferraz A, Herrero AI, et al. Anxiety, memory impairment, and locomotor dysfunction caused by a mutant thyroid hormone receptor alpha1 can be ameliorated by T3 treatment. Genes & development. Sep 15 2005;19(18):2152–63. doi:10.1101/gad.346105

36. Minakhina S, Bansal S, Zhang A, et al. A Direct Comparison of Thyroid Hormone Receptor Protein Levels in Mice Provides Unexpected Insights into Thyroid Hormone Action. Thyroid. Apr 6 2020;doi:10.1089/thy.2019.0763

37. Bernal J. Action of thyroid hormone in brain. J Endocrinol Invest. Mar 2002;25(3):268–88. doi:10.1007/BF03344003

38. Mayerl S, Müller J, Bauer R, et al. Transporters MCT8 and OATP1C1 maintain murine brain thyroid hormone homeostasis. J Clin Invest. May 2014;124(5):1987–99. doi:10.1172/jci70324

39. Yu L, Iwasaki T, Xu M, et al. Aberrant cerebellar development of transgenic mice expressing dominant-negative thyroid hormone receptor in cerebellar Purkinje cells. Endocrinology. Apr 2015;156(4):1565–76. doi:10.1210/en.2014-1079

40. Morte B, Manzano J, Scanlan T, Vennström B, Bernal J. Deletion of the thyroid hormone receptor alpha 1 prevents the structural alterations of the cerebellum induced by hypothyroidism. Proceedings of the National Academy of Sciences of the United States of America. Mar 19 2002;99(6):3985–9. doi:10.1073/pnas.062413299

41. Tinnikov A, Nordstrom K, Thoren P, et al. Retardation of post-natal development caused by a negatively acting thyroid hormone receptor alpha1. The EMBO journal. Oct 01 2002;21(19):5079–87.

42. Pelletier P, Gauthier K, Sideleva O, Samarut J, Silva JE. Mice lacking the thyroid hormone receptor-alpha gene spend more energy in thermogenesis, burn more fat, and are less sensitive to high-fat diet-induced obesity. Endocrinology. Dec 2008;149(12):6471–86. doi:10.1210/en.2008-0718

43. Sjogren M, Alkemade A, Mittag J, et al. Hypermetabolism in mice caused by the central action of an unliganded thyroid hormone receptor alpha1. The EMBO journal. Oct 31 2007;26(21):4535–45. doi:10.1038/sj.emboj.7601882

44. Jornayvaz FR, Lee HY, Jurczak MJ, et al. Thyroid hormone receptor-alpha gene knockout mice are protected from diet-induced hepatic insulin resistance. Endocrinology. Feb 2012;153(2):583–91. doi:10.1210/en.2011-1793

45. Eom YS, Wilson JR, Bernet VJ. Links between Thyroid Disorders and Glucose Homeostasis. Diabetes Metab J. Mar 2022;46(2):239–256. doi:10.4093/dmj.2022.0013

46. Verga Falzacappa C, Panacchia L, Bucci B, et al. 3,5,3’-triiodothyronine (T3) is a survival factor for pancreatic beta-cells undergoing apoptosis. Journal of cellular physiology. Feb 2006;206(2):309–21. doi:10.1002/jcp.20460

47. Verga Falzacappa C, Mangialardo C, Raffa S, et al. The thyroid hormone T3 improves function and survival of rat pancreatic islets during in vitro culture. Islets. Mar-Apr 2010;2(2):96–103. doi:10.4161/isl.2.2.11170

48. Furuya F, Shimura H, Yamashita S, Endo T, Kobayashi T. Liganded thyroid hormone receptor-alpha enhances proliferation of pancreatic beta-cells. J Biol Chem. Aug 6 2010;285(32):24477–86. doi:10.1074/jbc.M109.100222

49. Furuya F, Shimura H, Asami K, et al. Ligand-bound thyroid hormone receptor contributes to reprogramming of pancreatic acinar cells into insulin-producing cells. J Biol Chem. May 31 2013;288(22):16155–66. doi:10.1074/jbc.M112.438192

50. Verga Falzacappa C, Petrucci E, Patriarca V, et al. Thyroid hormone receptor TRbeta1 mediates Akt activation by T3 in pancreatic beta cells. J Mol Endocrinol. Feb 2007;38(1-2):221–33. doi:10.1677/jme.1.02166

51. Verga Falzacappa C, Patriarca V, Bucci B, et al. The TRbeta1 is essential in mediating T3 action on Akt pathway in human pancreatic insulinoma cells. Journal of cellular biochemistry. Apr 1 2009;106(5):835–48. doi:10.1002/jcb.22045

52. Blanchet E, Pessemesse L, Feillet-Coudray C, et al. p43, a Truncated Form of Thyroid Hormone Receptor α, Regulates Maturation of Pancreatic β Cells. International journal of molecular sciences. Mar 2 2021;22(5)doi:10.3390/ijms22052489

53. Lin Y, Sun Z. Thyroid hormone promotes insulin-induced glucose uptake by enhancing Akt phosphorylation and VAMP2 translocation in 3T3-L1 adipocytes. Journal of cellular physiology. Oct 2011;226(10):2625–32. doi:10.1002/jcp.22613

54. Lin Y, Sun Z. Thyroid hormone potentiates insulin signaling and attenuates hyperglycemia and insulin resistance in a mouse model of type 2 diabetes. British journal of pharmacology. Feb 2011;162(3):597–610. doi:10.1111/j.1476-5381.2010.01056.x

55. Weinstein SP, O’Boyle E, Haber RS. Thyroid hormone increases basal and insulin-stimulated glucose transport in skeletal muscle. The role of GLUT4 glucose transporter expression. Diabetes. Oct 1994;43(10):1185–9.

56. Teixeira SS, Tamrakar AK, Goulart-Silva F, Serrano-Nascimento C, Klip A, Nunes MT. Triiodothyronine acutely stimulates glucose transport into L6 muscle cells without increasing surface GLUT4, GLUT1, or GLUT3. Thyroid. Jul 2012;22(7):747–54. doi:10.1089/thy.2011.0422

57. Segal J, Ingbar SH. In vivo stimulation of sugar uptake in rat thymocytes. An extranuclear action of 3,5,3’-triiodothyronine. J Clin Invest. Oct 1985;76(4):1575–80. doi:10.1172/jci112139

58. Champy MF, Selloum M, Zeitler V, et al. Genetic background determines metabolic phenotypes in the mouse. Mamm Genome. May 2008;19(5):318–31. doi:10.1007/s00335-008-9107-z

59. Bahi L, Garnier A, Fortin D, et al. Differential effects of thyroid hormones on energy metabolism of rat slow- and fast-twitch muscles. Journal of cellular physiology. Jun 2005;203(3):589–98. doi:10.1002/jcp.20273

60. Mohamed F, Endre Z, Jayamanne S, et al. Mechanisms underlying early rapid increases in creatinine in paraquat poisoning. PloS one. 2015;10(3):e0122357. doi:10.1371/journal.pone.0122357

61. Basu G, Mohapatra A. Interactions between thyroid disorders and kidney disease. Indian J Endocrinol Metab. Mar 2012;16(2):204–13. doi:10.4103/2230-8210.93737

62. Patil VP, Shilpasree AS, Patil VS, Pravinchandra KR, Ingleshwar DG, Vani AC. Evaluation of renal function in subclinical hypothyroidism. J Lab Physicians. Jan-Mar 2018;10(1):50–55. doi:10.4103/jlp.Jlp_67_17

63. Maffei M, Halaas J, Ravussin E, et al. Leptin levels in human and rodent: measurement of plasma leptin and ob RNA in obese and weight-reduced subjects. Nature medicine. Nov 1995;1(11):1155–61. doi:10.1038/nm1195-1155

64. Oge A, Bayraktar F, Saygili F, Guney E, Demir S. TSH influences serum leptin levels independent of thyroid hormones in hypothyroid and hyperthyroid patients. Endocr J. Apr 2005;52(2):213–7. doi:10.1507/endocrj.52.213

65. Akbaba G, Berker D, Isık S, et al. Changes in the before and after thyroxine treatment levels of adipose tissue, leptin, and resistin in subclinical hypothyroid patients. Wien Klin Wochenschr. Aug 2016;128(15-16):579–85. doi:10.1007/s00508-015-0865-9

66. Ballas SK. Normal serum iron and elevated total iron-binding capacity in iron-deficiency states. Am J Clin Pathol. Apr 1979;71(4):401–3. doi:10.1093/ajcp/71.4.401

67. Wopereis DM, Du Puy RS, van Heemst D, et al. The relation between thyroid function and anemia: a pooled analysis of individual participant data. J Clin Endocrinol Metab. Aug 2 2018;doi:10.1210/jc.2018-00481

68. Lademann F, Tsourdi E, Rijntjes E, et al. Lack of the thyroid hormone transporter Mct8 in osteoblast and osteoclast progenitors both increases trabecular bone in male mice. Thyroid. Jan 5 2020;doi:10.1089/thy.2019.0271

69. Bremner AP, Feddema P, Joske DJ, et al. Significant association between thyroid hormones and erythrocyte indices in euthyroid subjects. Clinical endocrinology. Feb 2012;76(2):304–11. doi:10.1111/j.1365-2265.2011.04228.x

70. Kendrick TS, Payne CJ, Epis MR, et al. Erythroid defects in TRalpha-/- mice. Blood. Mar 15 2008;111(6):3245–8. doi:10.1182/blood-2007-07-101105

71. Angelin-Duclos C, Domenget C, Kolbus A, Beug H, Jurdic P, Samarut J. Thyroid hormone T3 acting through the thyroid hormone alpha receptor is necessary for implementation of erythropoiesis in the neonatal spleen environment in the mouse. Research Support, Non-U.S. Gov’t. Development. Mar 2005;132(5):925–34. doi:10.1242/dev.01648

72. Rotter D, Peiris H, Grinsfelder DB, et al. Regulator of Calcineurin 1 helps coordinate whole-body metabolism and thermogenesis. EMBO reports. Nov 2 2018;doi:10.15252/embr.201744706

73. van Gucht ALM, Meima ME, Moran C, et al. Anemia in Patients With Resistance to Thyroid Hormone alpha: A Role for Thyroid Hormone Receptor alpha in Human Erythropoiesis. J Clin Endocrinol Metab. Sep 01 2017;102(9):3517–3525. doi:10.1210/jc.2017-00840

74. Demir K, van Gucht AL, Büyükinan M, et al. Diverse Genotypes and Phenotypes of Three Novel Thyroid Hormone Receptor-α Mutations. J Clin Endocrinol Metab. Aug 2016;101(8):2945–54. doi:10.1210/jc.2016-1404

75. Jafari M, Ghadami E, Dadkhah T, Akhavan-Niaki H. PI3k/AKT signaling pathway: Erythropoiesis and beyond. Journal of cellular physiology. Mar 2019;234(3):2373–2385. doi:10.1002/jcp.27262

76. Rubingh J, van der Spek A, Fliers E, Boelen A. The Role of Thyroid Hormone in the Innate and Adaptive Immune Response during Infection. Comprehensive Physiology. Sep 24 2020;10(4):1277–1287. doi:10.1002/cphy.c200003

77. van der Spek AH, Fliers E, Boelen A. Thyroid Hormone and Deiodination in Innate Immune Cells. Endocrinology. Jan 1 2021;162(1)doi:10.1210/endocr/bqaa200

78. Wenzek C, Boelen A, Westendorf AM, Engel DR, Moeller LC, Führer D. The interplay of thyroid hormones and the immune system -where we stand and why we need to know about it. Eur J Endocrinol. Mar 23 2022;186(5):R65–r77. doi:10.1530/eje-21-1171

79. Arpin C, Pihlgren M, Fraichard A, et al. Effects of T3R alpha 1 and T3R alpha 2 gene deletion on T and B lymphocyte development. Journal of immunology (Baltimore, Md : 1950). Jan 1 2000;164(1):152–60. doi:10.4049/jimmunol.164.1.152

80. Aoki N, Wakisaka G, Nagata I. Effects of thyroxine on T-cell counts and tumour cell rejection in mice. Acta Endocrinol (Copenh). Jan 1976;81(1):104–9. doi:10.1530/acta.0.0810104

81. Frick LR, Rapanelli M, Bussmann UA, et al. Involvement of thyroid hormones in the alterations of T-cell immunity and tumor progression induced by chronic stress. Biol Psychiatry. Jun 1 2009;65(11):935–42. doi:10.1016/j.biopsych.2008.12.013

82. Billon C, Canaple L, Fleury S, et al. TRalpha Protects Against Atherosclerosis in Male Mice: Identification of a Novel Anti-Inflammatory Property for TRalpha in Mice. Endocrinology. Jul 2014;155(7):2735–45. doi:10.1210/en.2014-1098

83. Park S, Zhu X, Kim M, Zhao L, Cheng SY. Thyroid Hormone Receptor α1 Mutants Impair B Lymphocyte Development in a Mouse Model. Thyroid. Jun 2021;31(6):994–1002. doi:10.1089/thy.2019.0782

84. Bianco AC, Dumitrescu A, Gereben B, et al. Paradigms of Dynamic Control of Thyroid Hormone Signaling. Endocr Rev. Aug 1 2019;40(4):1000–1047. doi:10.1210/er.2018-00275

85. Hodin RA, Lazar MA, Chin WW. Differential and tissue-specific regulation of the multiple rat c-erbA messenger RNA species by thyroid hormone. J Clin Invest. Jan 1990;85(1):101–5. doi:10.1172/JCI114398

86. Zhu X, Cheng SY. New insights into regulation of lipid metabolism by thyroid hormone. Current opinion in endocrinology, diabetes, and obesity. Oct 2010;17(5):408–13. doi:10.1097/MED.0b013e32833d6d46

87. Chaves C, Bruinstroop E, Refetoff S, Yen PM, Anselmo J. Increased Hepatic Fat Content in Patients with Resistance to Thyroid Hormone Beta. Thyroid. Jul 2021;31(7):1127–1134. doi:10.1089/thy.2020.0651

88. Gauthier K, Flamant F. Nongenomic, TRbeta-Dependent, Thyroid Hormone Response Gets Genetic Support. Endocrinology. Sep 2014;155(9):3206–9. doi:10.1210/en.2014-1597

89. Klieverik LP, Sauerwein HP, Ackermans MT, Boelen A, Kalsbeek A, Fliers E. Effects of thyrotoxicosis and selective hepatic autonomic denervation on hepatic glucose metabolism in rats. American journal of physiology Endocrinology and metabolism. Mar 2008;294(3):E513–20. doi:10.1152/ajpendo.00659.2007

90. Mittag J, Lyons DJ, Sallstrom J, et al. Thyroid hormone is required for hypothalamic neurons regulating cardiovascular functions. J Clin Invest. Jan 2 2013;123(1):509–16. doi:10.1172/jci65252

