## Supplement for "Comparative phenotyping of mice reveals canonical and noncanonical physiological functions of TRα and TRβ"

**Supplemental information**

**Figure S1**

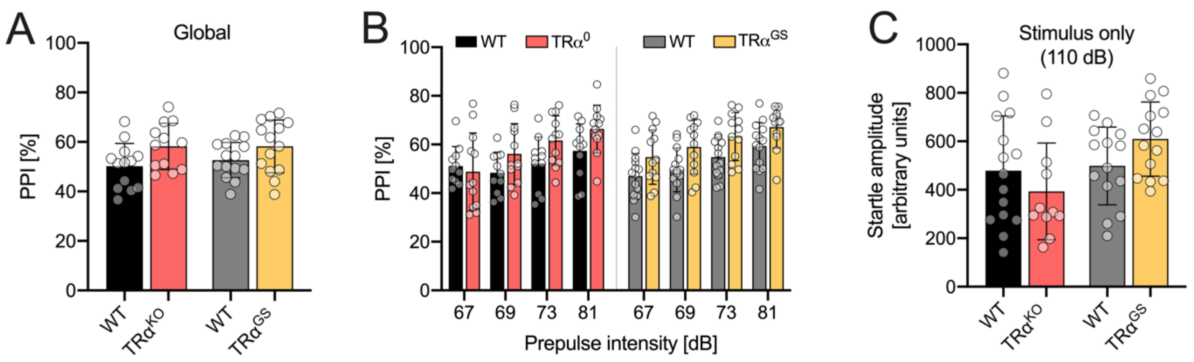


**Figure S1: Prepulse inhibition (PPI) of female TRα mutant mice. A** global PPI, calculated as the mean PPI [%] for the different prepulse responses. **B** PPI separated by prepulse intensities ranging from 67 to 81 dB. **C** Startle amplitude with a stimulus or 110 dB. (TRα, n=12-15).

**Figure S2**

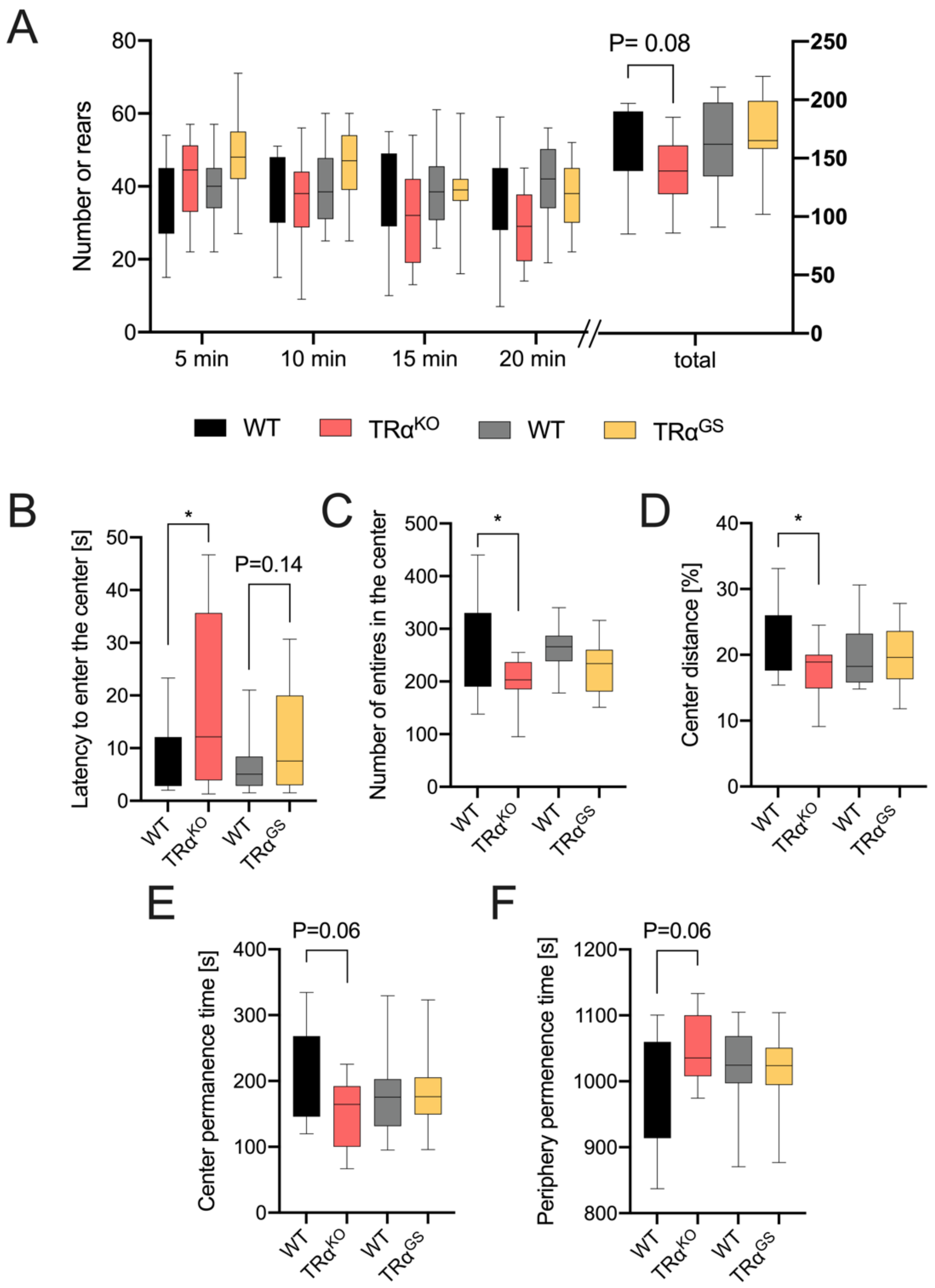


**Figure S2: Open field test of female TRα mice. A** Number of rears at certain time points and total number of rears (total) during 20 min test period. **B** Latency to enter the center. **C** Number of entries in the center. **D** Percentage of distance travelled in the center. **E** Time spent in the center and **F** time spent in the periphery. (n=11-15; One-way ANOVA with Sidak's multiple comparison test; *$P<0.05$).

**Figure S3**

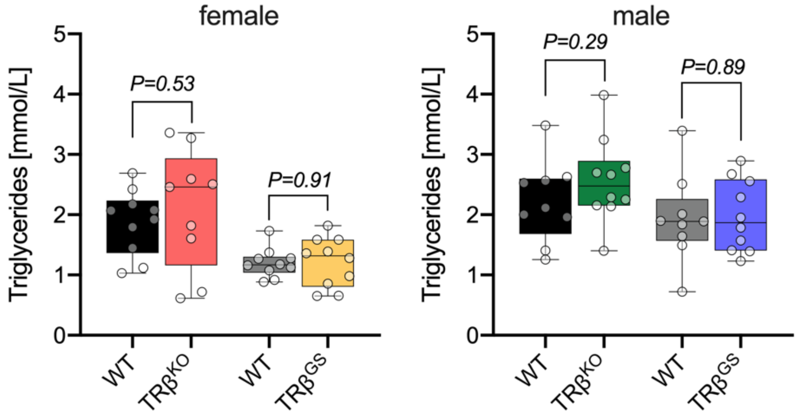


**Figure S3: Serum triglyceride content of female and male TRβ mouse models.** Triglyceride serum concentrations for female (left) and male (right) TRβ$^{KO}$ and TRβ$^{GS}$ mice compared to their WT controls. (One-way ANOVA; n=9-10).

### Figure S4

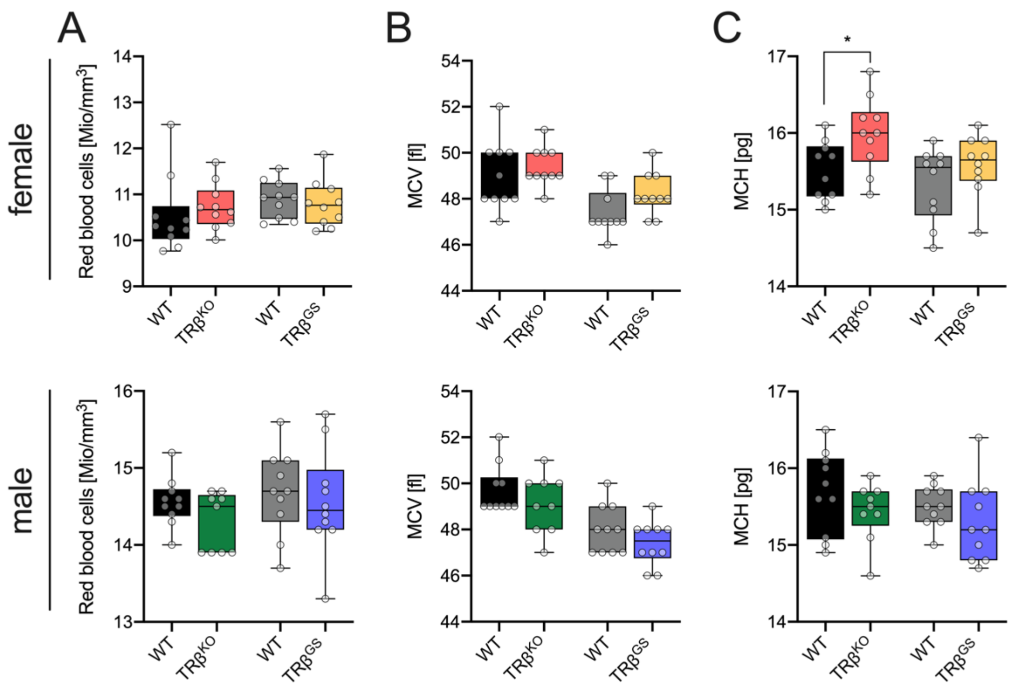


**Figure S4: Hematological analysis of TRβ mutant mice.** For both sexes of TRβ mouse lines red blood cell count (RBC) **A**, mean corpuscular volume (MCV) **B** and the mean corpuscular hemoglobin content (MCH) **C** were determined. One-way ANOVA; *$P<0.05$, (n=9-10).
